# Proof-of-concept of targeted degradation of p38α/β MAPK host-kinase as a potent inhibitor of coronaviruses

**DOI:** 10.64898/2026.04.29.721712

**Authors:** Gemma Cooper, Tim Snape, Maitreyi Shivkumar

**Affiliations:** Leicester School of Pharmacy

## Abstract

Host-targeting antivirals offer a promising strategy for combating emerging viral threats by targeting cellular pathways required for infection. The p38 mitogen-activated protein kinase (MAPK) pathway has been implicated as a host dependency factor exploited by multiple viruses, including coronaviruses, making it an attractive antiviral target. Here, we show for the first time that targeted degradation of p38 using the proteolysis-targeting chimera (PROTAC) NR-7h potently inhibits coronavirus infection. NR-7h induced substantial degradation of p38 in multiple cell lines and inhibited infection of two seasonal coronaviruses OC43 and 229E, providing broad pan-coronavirus activity. Infectious viral titres and viral RNA levels were significantly reduced without any detectable cytotoxicity. NR-7h exhibited greater antiviral potency than conventional p38 small-molecule inhibitors, with an IC_50_ of 1.0 nM compared with 648.4 nM for LY2228820, while the parent kinase inhibitor PH-797804 did not achieve 50% inhibition at the highest concentration tested. Pseudovirus and time-of-addition studies indicated that antiviral activity occurred at a post-entry stage of infection. Importantly, antiviral activity was eliminated by inhibition of proteasome function or E3 ligase activity, demonstrating dependence on PROTAC-mediated degradation. Our findings provide a proof-of-concept that targeted degradation of host kinase p38 can function as an antiviral modality and suggest PROTAC-based host-directed therapeutics may offer advantages over conventional kinase inhibition for broad-spectrum antiviral development.

## Introduction

The emergence of SARS-CoV-2 in 2019 resulted in the COVID-19 pandemic, causing millions of deaths worldwide and placing unprecedented strain on healthcare systems^1^. While vaccines have significantly reduced severe disease and mortality, the development and deployment of vaccines often lag behind viral emergence and variant evolution. In addition, most antiviral drugs are virus-specific, limiting their usefulness against newly emerging pathogens^2^. This highlights the need for antiviral strategies that can be rapidly deployed against current and future viral threats.

One promising approach is the development of host-targeting antivirals. Because viruses rely heavily on host cell machinery to complete their replication cycle, targeting cellular pathways required for viral infection may provide broader antiviral activity while reducing the likelihood of resistance mutations^3^. Host kinases have emerged as important regulators of virus-host interactions, controlling signalling pathways that viruses exploit to support entry, replication, and immune evasion ^4,5^.

The p38 mitogen-activated protein kinase (MAPK) signalling pathway plays a central role in cellular stress responses, inflammation, and immune signalling^6^. Increasing evidence suggests that viruses exploit this pathway to support replication and promote pathogenic effects^7–13^. Activation of p38 MAPK occurs through both canonical and non-canonical mechanisms. In the classical pathway, upstream kinases MKK3 and MKK6 phosphorylate p38 MAPK at threonine 180 and tyrosine 182, resulting in activation of the kinase^14^. Viral infection can trigger this cascade through cellular stress responses or receptor-mediated signalling events that occur during viral entry^15,16^. Coronaviruses have been shown to stimulate phosphorylation of MKK3/6, leading to increased p38 signalling and downstream activation of inflammatory and translational pathways^6,12^.

In addition to classical activation, p38 MAPK can also be activated through atypical mechanisms that bypass upstream kinases. In this pathway, the scaffold protein TAB1 binds directly to p38α, inducing autophosphorylation and activation of the kinase^17,18^. This atypical pathway is often more rapid and transient, typically occurring within minutes following stimulation. Some viruses may exploit this mechanism to rapidly activate p38 signalling and alter host cell function early during infection^19^.

Coronaviruses have evolved several strategies to activate the p38 MAPK pathway^7,17,20^. Viral entry through ACE2 has been shown to trigger excessive p38 activation, and disruption of ACE2 signalling during infection can increase angiotensin II levels and further enhance p38 activity^21^. In addition, several coronavirus proteins have been reported to stimulate p38 signalling directly. The SARS-CoV 3a protein has been associated with increased p53 expression and activation of p38^22,23^, while the nucleocapsid protein, papain-like protease, and 7a protein have also been linked to p38 activation^24–26^. Through these mechanisms, coronaviruses can drive sustained p38 signalling, creating a cellular environment that favours viral replication and inflammation.

Activation of p38 MAPK has also been shown to enhance phosphorylation of eukaryotic initiation factor 4E, a key regulator of cap-dependent translation^17,27^. This modification can increase the efficiency of protein synthesis, including translation of viral proteins required for replication^28^. In addition, prolonged p38 activation can promote apoptosis and inflammatory signalling, processes that may contribute to viral dissemination and disease severity^29^.

Consistent with this role in viral infection, inhibition of p38 MAPK signalling has been shown to impair replication of several viruses^30,31^, supporting its potential as a host-directed antiviral target. In human coronavirus 229E infection, small molecule inhibition of p38 reduced viral RNA levels by approximately 50% at 4-10 μM^30^, while in respiratory syncytial virus (RSV) infection p38 inhibition reduced progeny virus titres by 92-97% at 0.1-5 μM^32^. Similarly, the p38 inhibitor ralimetinib inhibited SARS-CoV-2 replication with an IC_50_ of 0.873 μM, further demonstrating antiviral activity through pharmacological targeting of this pathway^5^. The p38 inhibitor pamapimod has also shown anti-SARS-CoV-2 activity *in vitro* and progressed into clinical evaluation in combination with an anti-inflammatory agent, although the trial was later discontinued due to lower than expected efficacy in humans^33,34^. While these studies highlight the promise of p38 inhibition as an antiviral strategy, conventional small molecule kinase inhibitors can face challenges including poor oral bioavailability, driven by low aqueous solubility, high lipophilicity and extensive first-pass metabolism^35^.

Proteolysis-targeting chimeras (PROTACs) represent an emerging strategy for modulating protein function through targeted degradation rather than inhibition alone. PROTACs are bifunctional molecules consisting of a ligand that binds a protein of interest and a second ligand that recruits an E3 ubiquitin ligase, connected by a flexible linker^36^. By bringing the target protein into proximity with an E3 ligase, PROTACs promote ubiquitination of the target protein and subsequent degradation by the proteasome^37^ (Figure 1a). Unlike traditional small-molecule inhibitors, which require sustained occupancy of the target protein, PROTACs eliminate the protein entirely and can function catalytically within the cell^38^. Although PROTAC technology has primarily been developed in oncology, its potential application in antiviral therapy is gaining increasing attention. Both host-targeting degraders, such as cyclophilin A-targeting PROTACs with activity against HCV and HIV-1, and virus-targeting degraders directed against the SARS-CoV-2 main protease (Mpro) have been demonstrated as having as antiviral efficacies^39,40^. By enabling the selective degradation of host proteins required for viral infection, PROTACs may provide a strategy to disrupt virus-host interactions while reducing the likelihood of resistance^38,41^.

**Figure 1:**
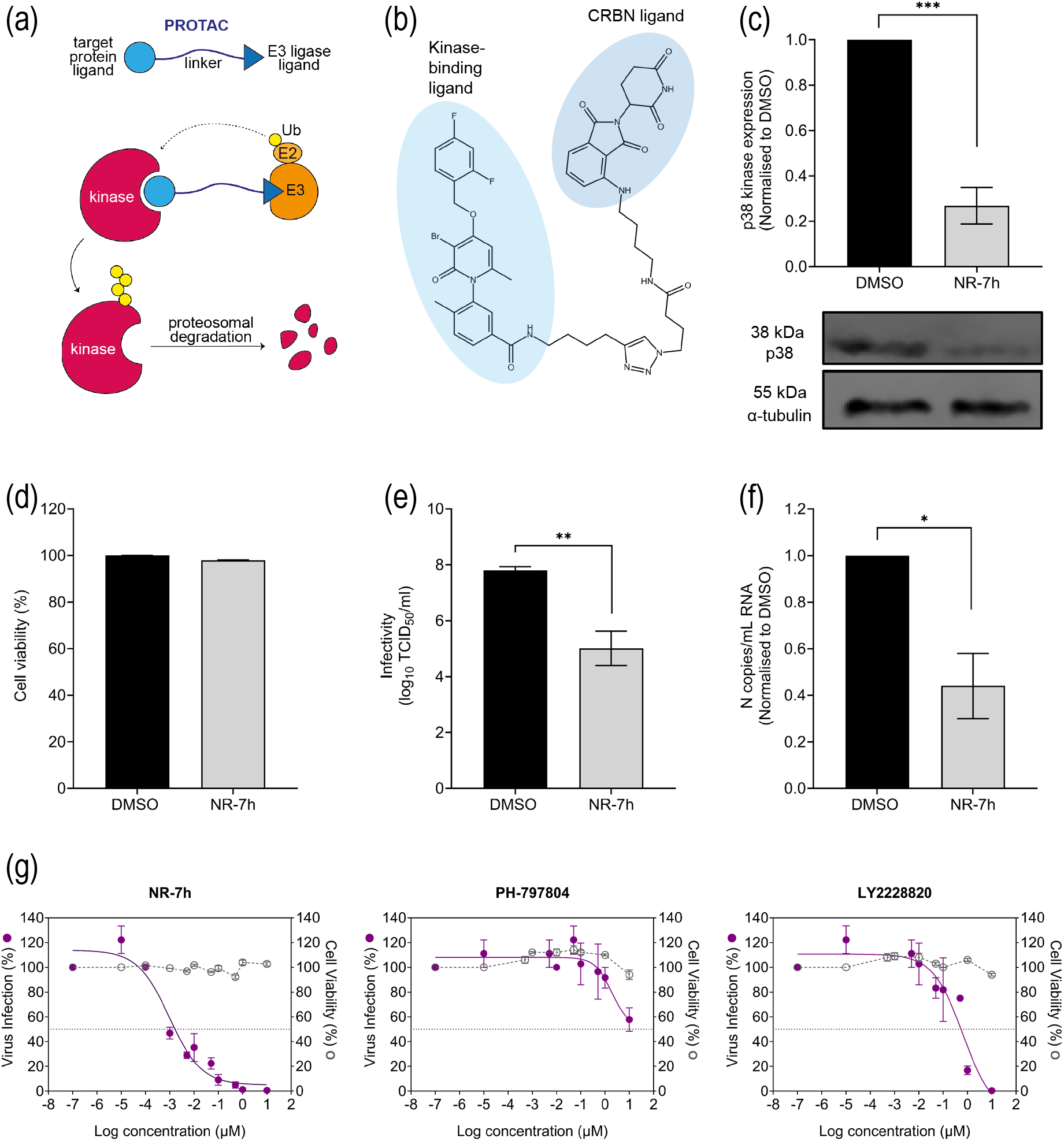
p38α/β-degrading PROTAC inhibits seasonal coronavirus OC43 in A549 cells. **(a)** PROTACs are bifunctional molecules, consisting of a target protein-binding ligand and an E3 ligase ligand attached via a linker. Upon binding of both targets, the E3 ligase catalyses ubiquitination of the target protein, inducing its degradation. **(b)** Structure of p38α/β-targeting PROTAC NR-7h. The shaded regions indicate the kinase-binding ligand (based on PH-797804) and the CRBN E3 ligase ligand. A549 cells treated with 10 μM NR-7h for 24 h show significant degradation of p38 kinase **(c)** but no effects on cell viability, as measured by MTS assay **(d)**. A549 cells infected with OC43 (MOI=0.1) and treated with 10 μM NR-7h showed significant reduction of viral infectivity **(e)** and viral RNA **(f)** after 72 h. **(g)** NR-7h showed greater antiviral efficacy compared to two p38 kinase small molecule inhibitors (NR-7h, IC_50_ 1.0 nM; PH-797804, IC_50_ >10 µM; LY2228820, IC_50_ 648.4 nM). Graphs show mean ± SEM (n=3); *, p≤0.05; **, p≤0.01; ***, p≤0.001.

We investigated the antiviral potential of NR-7h, a p38-targeting PROTAC (Figure 1b). NR-7h is a cereblon (CRBN)-recruiting PROTAC that selectively induces degradation of the p38α and p38β isoforms, without activity against other MAPKs, demonstrating high target specificity. It exhibits potent degradation with 50% degradation concentration (DC_50_) values in the nM range across multiple cell types^42^. Importantly, NR-7h does not alter p38 mRNA levels, indicating its effects are mediated through targeted protein degradation rather than transcriptional regulation^42^. NR-7h has also shown antiviral activity against the alphavirus Mayaro virus in primary human dermal fibroblasts and HeLa cells, supporting the potential of p38-targeting PROTACs as host-directed antivirals^11^.

In this study, we show that NR-7h has antiviral activity against coronaviruses, successfully inhibiting seasonal coronaviruses OC43 and 229E in a range of cell lines. The antiviral activity is proteosome-dependent, and the PROTAC shows higher efficacies compared to the corresponding p38α/β small-molecule inhibitor. We provide a proof-of-concept for using PROTACs that selectively degrade host kinases as pan-coronavirus inhibitors.

## Results

We first assessed the action of p38α/β-targeting PROTAC, NR-7h, against seasonal human coronavirus OC43 (Figure 1). To confirm target engagement in our infection models, p38 degradation was determined following treatment with NR-7h in human lung A549 cells. Treatment with 10 µM NR-7h for 24 h reduced p38 protein levels by 74% compared to untreated controls (Figure 1c), with no impact on cell viability (Figure 1d). Next, A549 cells infected with OC43 were treated with 10 µM NR-7h for 72 h, after which viral infectivity was assessed. OC43 titres showed a 2.8 log_10_ reduction compared to the DMSO-treated controls, indicating potent antiviral activity (Figure 1e). Consistent with this, viral RNA levels in culture supernatants were also significantly decreased following treatment (Figure 1f). These results were also recapitulated in BHK-21 epithelial cells, which showed significant reduction in OC43 infection and viral RNA with no effect on cell viability (Figure S1).

To compare targeted degradation with conventional kinase inhibition by small molecule inhibitors, antiviral activity of NR-7h was compared to two p38α/β inhibitors, PH-797804 and LY2228820 (Figure 1g). Neither NR-7h nor the small molecule inhibitors showed cytotoxicity in A549 cells up to 10 µM. NR-7h exhibited greater antiviral potency than either of the small molecule inhibitors, with an IC_50_ of 1.0 nM. LY2228820 showed an IC_50_ of 648.4 nM, while PH-797804, the parent kinase-binding ligand of NR-7h, did not reach 50% inhibition at the highest concentration tested of 10 µM. Thus, targeted degradation outperformed inhibition of p38α/β under the same conditions.

### Antiviral activity occurs at a post-entry stage

To determine if targeting p38 affects viral entry, we tested NR-7h in spike pseudovirus infection assays (Figure 2). HeLa-ACE2 cells were infected with SARS-CoV-1 or SARS-CoV-2 spike-pseudotyped lentiviruses expressing a luciferase reporter and treated with 10 µM NR-7h. The spike pseudoviruses recapitulate the ACE2-mediated entry mechanism of full-length viruses^43^, and therefore provide a model to investigate binding, entry and internalisation of coronaviruses. NR-7h treatment showed significant depletion of p38 (Figure 2a), while no cytotoxicity was observed in HeLa-ACE2 cells (Figure 2b). PROTAC treatment did not reduce pseudovirus infection relative to untreated controls, indicating no inhibition of spike-mediated entry (Figure 2c,d). This suggests that the antiviral effect of p38 degradation occurs at a post-entry stage of the viral life cycle.

**Figure 2:**
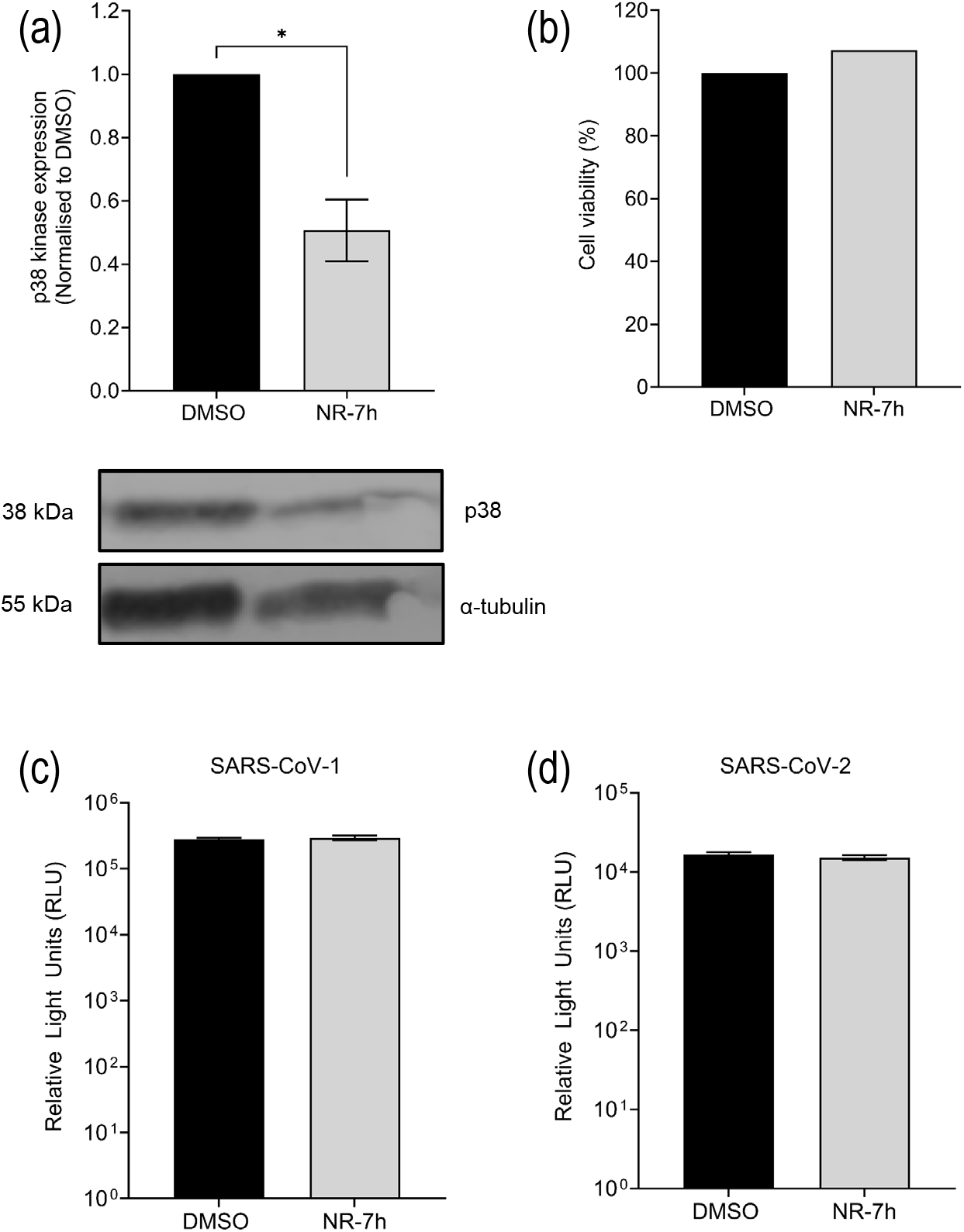
p38α/β-degrading PROTAC does not inhibit viral entry. HeLa-ACE2 cells treated with 10 μM NR-7h for 24 h show significant degradation of p38 kinase **(a)** but no effects on cell viability, as measured by MTS assay **(b)**. HeLa-ACE2 cells infected with SARS-CoV-1 **(c)** and SARS-CoV-2 **(d)** spike pseudoviruses and treated with 10 μM NR-7h showed no effect on viral infectivity after 72 h. Graphs show mean ± SEM (n=3); *, p≤0.05.

To define the stage of the viral life cycle affected by p38 degradation, time-of-addition experiments were performed in BHK-21 cells (Figure S2), in which 10 µM NR-7h was added either during infection or at 1, 3 or 6 h post-infection, and infection was quantified by immunofluorescence 14 h later. In contrast to the robust antiviral activity observed during longer treatment periods, no statistically significant inhibition was observed at these early infection time points, suggesting that p38 degradation is insufficient within this shorter window to affect a single infection cycle. This was consistent with our observations that ∼50% of p38 is degraded in BHK-21 cells as early as 3 h after treatment, however, p38 levels continued to decrease to less than 20% of untreated controls by 72 h (Figure S3). This suggests that antiviral activity requires longer exposure to NR-7h to establish sufficient target depletion prior to or during multi-cycle virus replication.

### Antiviral activity is dependent on E3 ligase recruitment and proteasomal degradation

To confirm that the antiviral activity of NR-7h was due to PROTAC-mediated degradation of p38, the requirement for E3 ligase activity and proteasomal function was assessed using MLN4924 and MG132 (Figure 3). MG132 is a proteasome inhibitor that blocks degradation of ubiquitinated proteins^44^. MLN4924 inhibits NEDD8-activating enzyme (NAE) function, preventing cullin neddylation and thereby inactivating Cullin-RING ligase (CRL) complexes, including the CRL4-CRBN ligase recruited by NR-7h^45^.

**Figure 3:**
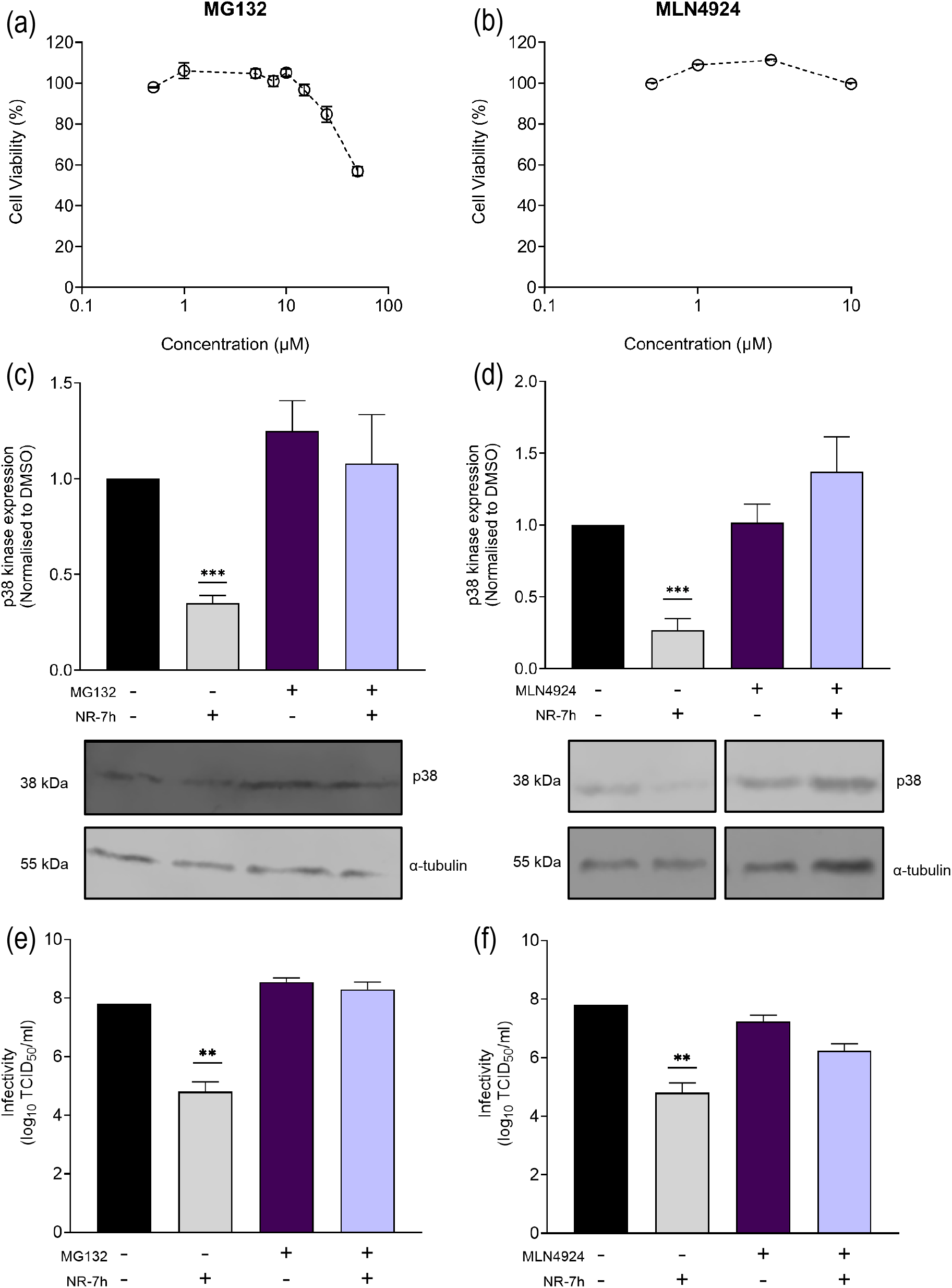
Antiviral activity of p38α/β-degrading PROTAC is proteosome-dependent and requires E3 ligase recruitment. **Cell viability of A549 cells treated with (a) MG132 or (b) MLN4924, no cytotoxicity was observed up to 10** µM for both inhibitors. A549 cells treated with 10 µM NR-7h and **(c)** MG132 (5 µM) or **(d)** MLN4924 (3 µM) showed no degradation of p38 kinase compared to when no inhibitors were present. A549 cells infected with OC43 (MOI=0.1) showed significant reduction in infection 72 h after treatment in presence of NR-7h, but infectivity was recovered in presence of **(e)** MG132 (5 µM) or **(f)** MLN4924 (3 µM). Graphs show mean ± SEM (n=3); **, p≤0.01; ***, p≤0.001.

To identify a suitable non-toxic concentration for the infection assays, MG132 cytotoxicity was first assessed in A549 cells over 72 h. Concentrations up to 5 µM were well-tolerated, while higher concentrations showed toxicity (Figure 3a). Western blot analysis demonstrated that 5 µM MG132 was sufficient to prevent NR7h-mediated degradation of p38, and co-treatment with MG132 inhibited target degradation compared to cells treated with NR-7h alone (Figure 3c). Importantly, co-treatment with 5 µM MG132 also reversed the antiviral activity of NR-7h during OC43 infection (Figure 3e). These data confirm that proteasome activity is required for both target degradation and antiviral efficacy.

MLN4924 exhibited low toxicity in A549 cells, with concentrations up to 10 µM showing no cytotoxicity (Figure 3b). Inhibition of NAE activity with MLN4924 similarly impaired NR-7h-mediated p38 degradation and reduced antiviral activity, confirming dependence on CRL-mediated ubiquitination and E3 ligase recruitment (Figure 3d,f).

These results demonstrate that both target degradation and antiviral activity require an intact ubiquitin–proteasome pathway, and indicate that the antiviral effects of NR-7h arise likely through PROTAC-mediated degradation rather than off-target compound activity.

### NR-7h exhibits pan-coronavirus antiviral activity

To determine if antiviral activity extended beyond OC43, NR-7h was evaluated against a seasonal alphacoronavirus 229E in MRC-5 cells (Figure 4). Western blot analysis confirmed up to 50% degradation of p38 in MRC-5 cells following 24 h treatment (Figure 4a). Consistent with results observed for OC43, treatment with 10 µM NR-7h significantly reduced infectious viral titres (3.1 log_10_ reduction) compared with DMSO-treated controls (Figure 4c), with no detectable effect on cell viability (Figure 4b). Viral RNA levels were also significantly reduced following treatment, indicating impaired viral replication (Figure 4d).

**Figure 4:**
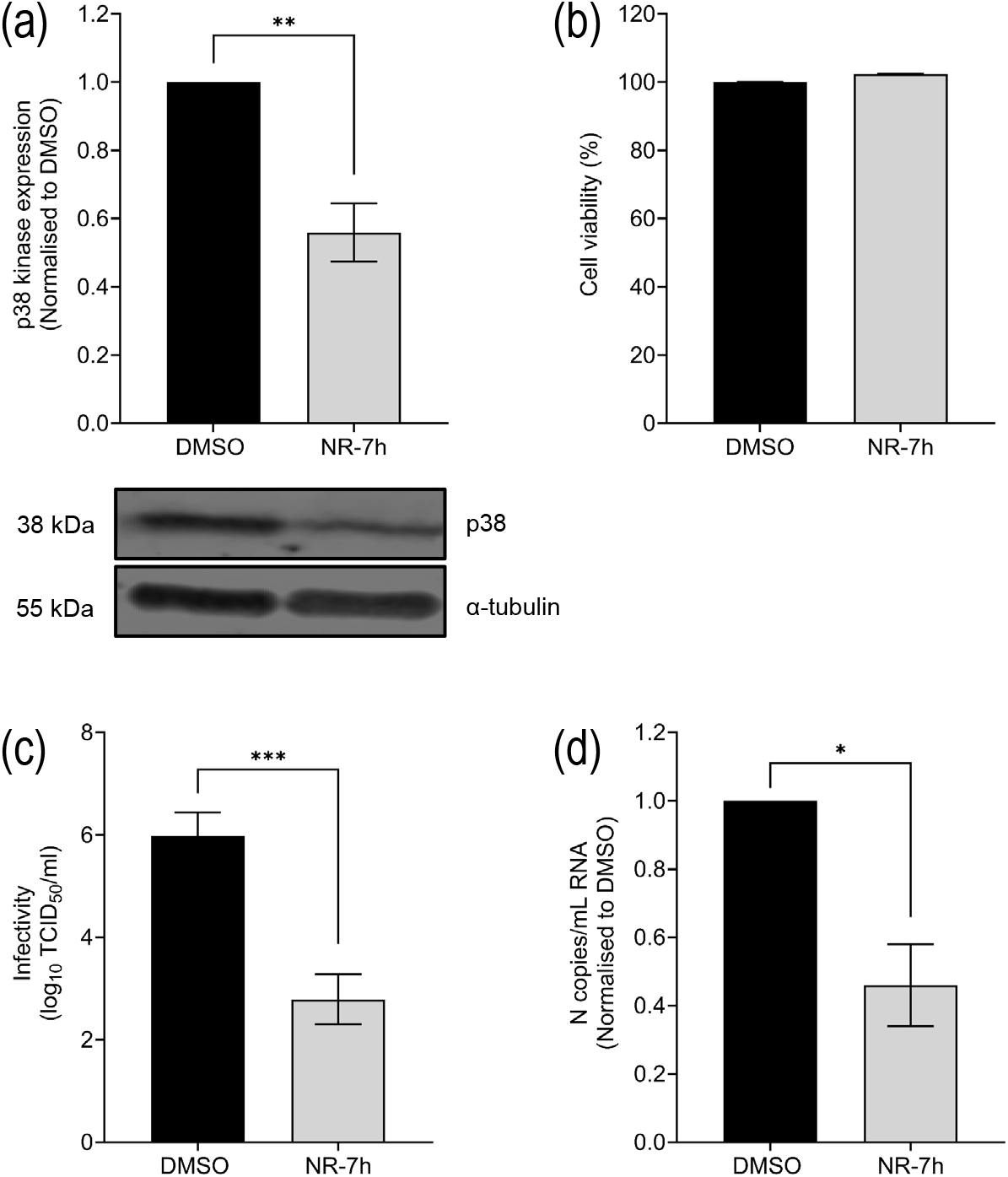
p38α/β-degrading PROTAC inhibits seasonal coronavirus 229E. MRC-5 cells treated with 10 μM NR-7h for 24 h show significant degradation of p38 kinase **(a)** but no effects on cell viability, as measured by MTS assay **(b)**. MRC-5 cells infected with 229E (MOI=0.1) and treated with 10 μM NR-7h showed significant reduction of viral infectivity **(c)** and viral RNA **(d)** after 72 h. Graphs show mean ± SEM (n=3); *, p≤0.05; **, p≤0.01; ***, p≤0.001.

Together, these findings support a role for p38 as a host factor exploited during coronavirus replication. The antiviral activity observed against both OC43 and 229E demonstrate that targeted degradation of p38 exerts pan-coronavirus activity against seasonal coronaviruses.

## Discussion

We demonstrate that targeted protein degradation using proteolysis-targeting chimeras (PROTACs) against the host kinase p38 MAPK can significantly inhibit infection by two seasonal coronaviruses. By selectively degrading this host dependency factor, PROTAC treatment reduced viral infectivity and viral RNA levels in infected cells without causing cytotoxicity at effective concentrations. The antiviral activity correlated with the degree of target protein degradation, suggesting that depletion of the kinase from the cellular environment is sufficient to disrupt viral replication. These findings support the concept that host-directed protein degradation represents a viable antiviral strategy and highlight the potential of targeting the p38 MAPK pathway to suppress coronavirus infection.

Compared with conventional small-molecule inhibitors, PROTAC-mediated degradation may offer advantages in antiviral potency at lower concentrations. The strong activity observed for NR-7h, with an IC_50_ of 1.0 nM, contrasts with the higher inhibitory concentrations observed for the small-molecule inhibitors LY2228820 (0.6 μM) and PH-797804, which did not achieve 50% inhibition at concentrations up to 10 μM. This is further supported by previously reported antiviral activity of LY2228820 (ralimetinib) against SARS-CoV-2 (IC_50_ 0.873 μM), and for another p38 inhibitor pamapimod (100-250 nM depending on cell type), both of which also exhibit higher IC_50_ values than NR-7h^5,33^. Similar observations have been made with other antiviral degraders, including a cyclophilin A-targeting PROTAC reported to have greater anti-HIV-1 activity than its parent small-molecule inhibitor, particularly at lower concentrations^46^. Together, these findings support the idea that degradation-based approaches may provide advantages over traditional occupancy-driven inhibitors, potentially through the catalytic and sustained nature of target removal.

PROTAC technology offers several potential advantages over traditional small-molecule inhibition. Unlike conventional inhibitors, which block catalytic activity but leave the protein intact, PROTACs recruit the cellular ubiquitin–proteasome system to induce degradation of the target protein^38^. This mechanism removes both catalytic and non-catalytic functions of the protein and may therefore produce more complete disruption of signalling pathways exploited by viruses.

The catalytic nature of PROTAC-mediated degradation may also contribute to enhanced efficacy. Because PROTAC molecules can induce degradation of multiple target proteins sequentially, they can produce sustained suppression of signalling pathways even at relatively low concentrations^47^. This property may provide advantages over conventional inhibitors that require continuous occupancy of the target protein to maintain activity.

Importantly, the antiviral activity was dependent on proteasome-mediated degradation. NR-7h alone reduced p38 protein levels by approximately 60%, whereas no reduction was observed in the presence of either MG132 or the NAE inhibitor MLN4924. This inhibition of p38 degradation was accompanied by a reduction in antiviral activity during OC43 infection, with virus infectivity recovering to untreated levels in presence of MG132 or MLN4924 alongside NR-7h. Disruption of either proteasome function or CRL E3 ligase activation was sufficient to prevent both target degradation and antiviral activity. This indicates that NR-7h activity depends on effective removal of the target protein rather than off-target chemical effects and supports a requirement for CRL-dependent ubiquitination and downstream proteasomal processing for the observed antiviral activity.

The results of this study support the growing body of evidence indicating that host-directed antiviral strategies can provide an effective alternative to traditional virus-targeting therapeutics^48–53^. Host-targeting antiviral strategies offer several potential advantages compared with direct-acting antivirals. Because host proteins mutate less frequently than viral proteins, therapies targeting host pathways may reduce the likelihood of resistance emerging during treatment ^49,54^. In addition, many viruses rely on common cellular pathways to support their replication, raising the possibility that host-directed therapies could exhibit broad-spectrum antiviral activity against multiple viral pathogens.

However, targeting host proteins also presents potential challenges. Many host signalling pathways play essential roles in normal cellular physiology, and therefore inhibition or degradation of these proteins may produce unwanted toxicity^55,56^. In this study, cytotoxicity was generally low at antiviral concentrations, suggesting that selective targeting of p38 MAPK can be tolerated within a therapeutic window. Nevertheless, further studies in additional cell types and *in vivo* models will be required to fully evaluate the safety profile of these compounds.

Another consideration is the potential for off-target effects associated with both targeting kinases and using PROTACs. Selective degradation can be achieved with rational PROTAC design. For instance, NR-7h has been shown to selectively degrade the p38α and p38β isoforms at nM concentrations, without affecting other MAPK family members^42^. The p38 isoforms also differ in their expression patterns, with p38α and p38β being widely expressed across tissues, while p38γ and p38δ show more restricted, tissue-specific expression^57^. Further, although PROTACs are designed to selectively recruit specific target proteins to E3 ligases, unintended interactions may occur depending on linker composition and ligand binding specificity. Indeed, the structural similarity between thalidomide and the CRBN-ligand has been suggested to potentially lead to risks of teratogenicity^58^, therefore alternative E3 ligases could be preferable. Careful optimisation of PROTAC design will therefore be important for future studies.

Overall, the results of this study highlight the potential of p38-targeted protein degradation as a successful strategy for antiviral therapy. By exploiting the cellular ubiquitin–proteasome system to remove host proteins required for viral replication, PROTAC technology provides a promising platform for disrupting virus–host interactions effectively. Future work should focus on optimising p38-targeting PROTACs for improved potency, selectivity, and pharmacological properties, as well as evaluating their antiviral activity in additional cellular and *in vivo* models.

## Materials and Methods

### Cell lines and viruses

All cell lines were cultured at 37 °C with 5% CO_2_. HCT-8 cells (ATCC CCL-244) were cultured in RPMI 1640 medium (Lonza) supplemented with 10% foetal bovine serum (FBS; HyClone) and 1% penicillin-streptomycin (100 IU/mL penicillin, 100 μg/mL streptomycin; Lonza). Baby hamster kidney 21 (BHK-21; ATCC CCL-10) cells, HeLa-ACE2 cells (kind gift from James E Voss, TSRI, USA^59^), human lung epithelial cells (MRC-5; EACC #05072101), and human embryonic kidney (HEK) 293T cells were cultured in Dulbecco’s Modified Eagle Medium (DMEM, Lonza) containing 10% FBS and 1% penicillin-streptomycin.

Human coronaviruses OC43 (ATCC VR-1558) stocks were grown in HCT-8 cells, and human coronavirus 229E (ATCC VR-740) stocks were grown and titred in MRC-5 cells. Cells were infected with virus (MOI = 0.1) in media supplemented with 5% FBS, and incubated at 33 °C, 5% CO_2_ for 7 days. Cells were freeze/thawed at -80 °C three times, and cells and supernatant containing virus pooled, and centrifuged (1200 x*g*, 4 min) to remove cell debris. Supernatant was sterile-filtered (0.2 μm PES filter), and aliquots stored at -80 °C.

### Cell viability assay

Cell viability was assessed by a colorimetric MTS assay. Cell monolayers were treated with compound or inhibitor dilutions, and incubated at 33 °C, 5% CO_2_ for three days. MTS tetrazolium reagent (Abcam, ab223881) was mixed with phenazine methosulfate (PMS; Sigma, P9625) at a 20:1 ratio (for final concentrations of 2 mg/mL MTS and 46 μg/mL PMS). 20 μL MTS/PMS mixture per 100 μL media was added to the cells and plate incubated at 37 °C for 1 h in the dark. Absorbance at 490 nm was measured (SpectraMAX 5, Molecular Devices), and cell viability normalised to the DMSO controls.

### Compound treatment

NR-7h (Tocris Bioscience, #7177), PH-797804 (Stratech, A8308-APE), LY2228820 (Stratech, A5566-APE), MG132 (Sigma, # 474790) and MLN4924 (MedChemExpress, HY-70062) stocks were prepared in dimethyl sulfoxide (DMSO) and filtered through a 0.2 μm PES filter. Aliquots were stored at -20 °C.

A549 (OC43) or MRC-5 (229E) cells were seeded 24 h prior to treatment, and compounds or inhibitors diluted to required concentrations in infection media (DMEM supplemented with 5% FBS). Cells were infected with OC43 or 229E at MOI 0.1 for 1 h, compounds added, and incubated at 33 °C, 5% CO_2_ for 3 days. The virus in supernatants was quantified by a 50% tissue culture dose (TCID_50_) assay.

### Virus infectivity quantification

Virus stocks and viral titres following treatments were quantified by 50% tissue culture dose (TCID_50_) assays. Serial dilutions of virus stocks or treatment supernatants in infection media (DMEM supplemented with 5% FBS) were transferred to BHK-21 (OC43) or MRC-5 (229E) cell monolayers in 96-well plates, and incubated at 33 °C, 5% CO_2_. After 4 days, plates were scored for cytopathic effect (CPE) and TCID_50_ calculated using the Karber method (Lambert et al. 2008). The limit of detection for the assay was 0.80 log_10_ TCID_50_/mL.

### Viral RNA quantification

Following treatment with NR-7h, viral RNA was extracted from infected cells using the Monarch RNA extraction kit (NEB) according to manufacturer’s instructions. RNA was eluted in 100 μL nuclease free water and stored at -80 °C until use.

Reverse transcription quantitative PCR (RT-qPCR) was conducted to quantify viral RNA, using the SuperScriptTM III One-Step RT-PCR System with Platinum Taq DNA polymerase (Thermo Fisher) according to manufacturer’s instructions. OC43 N primers and probe (forward, 5’-AGCAACCAGGCTGATGTCAATACC-3’; reverse, 5’- AGCAGACCTTCCTGAGCCTTCAAT’-3; probe, 5’-[6FAM] TGACATTGTCGATCGGGACCCAAGTA [TAM]-3’)^60^ and 229E N primers and probe (forward, 5’-TCTCTTTATAGCCCTTTGCTTG-3’; reverse, 5’-ACCCGTTTGCCCTTTCTAGT-3’; probe, 5’-[6FAM] CAACCTTGGAAGGTGATACCTCGT [TAM]-3’)^61^ were used. RT-qPCR was carried out in QuantStudio 5 real-time PCR system (Applied Biosystems), using the following parameters: 50 °C for 15 min hold; 95 °C for 2 min; 50 cycles of 95 °C for 15 s, 60 °C for 30 s. Amount of viral RNA was normalised by comparing to a standard curve of known quantity of infectious virus (8.0 log_10_ TCID_50_/mL OC43 or 6.5 log_10_ TCID_50_/mL 229E). Samples were run in technical duplicates, and the standard curve was run in triplicate.

### Pseudovirus assays

Pseudoviruses were produced by transfecting HEK 293T cells with 1.3 μg p8.91 (HIV-1 gag-pol expression vector), 1.3 μg pCSFLW (containing a luciferase reporter), and 200 ng of plasmid expressing SARS-CoV-1 or SARS-CoV-2 spike protein, using FuGENE 6 transfection reagent (Promega). Cells were incubated at 37 °C for 48 h, after which pseudovirus supernatant was collected through a 0.2 μm PES filter.

HeLa-ACE2 cells were infected with pseudovirus and treated with NR-7h (or DMSO control) and incubated at 37 °C for 3 days. Infectivity was quantified by using the BrightGlo luciferase assay system (Promega).

### Time of addition assay

For time of addition experiments, cell monolayers were infected with virus for 1 h, and incubated in media until replacing with compounds at 0, 1, 3, or 6 h post-infection. Cells were incubated at 33 °C for 14 h, after which infectivity was quantified by immunofluorescence.

Cells were fixed in ice-cold methanol at 4 °C for 20 min, then permeabilised in 1% Triton-X 100 for 15 min, and washed in PBS. Cells were stained with OC43 N-specific primary antibody (Sigma, MAB9013, 1:2000) for 1 h and a goat anti-mouse secondary antibody (Invitrogen, F2761, 1:1000) for 1 h, and finally with 300 nM DAPI (Sigma, D9542) for 10 min. Plates were imaged and analysed on a High-Content Screening System (Cellomics).

### Quantification of protein degradation

Cells were seeded 24 h prior to treatment. After the specified treatment time, cells were washed in ice-cold PBS, detached using Trypsin-EDTA (Sigma), washed again in ice-cold PBS, and lysed in RIPA buffer (5 mM Tris-HCl pH 8, 150 mM NaCl, 1% Triton X-100, 1% sodium deoxycholate, 0.1% SDS). Lysates were centrifuged (17,000 x*g*, 10 min, 4 °C). Protein-containing supernatants were stored at -80 °C, until quantification by Western blotting.

Cell lysates were reduced in Laemmli buffer (60 mM Tris-HCl pH 6.8, 2% SDS, 10% glycerol, 5% β-mercaptoethanol, 0.01% bromophenol blue), denatured by incubating at 95 °C for 5 min, then cooled on ice. Samples and protein ladder were run on a 10% Bis-Tris protein gel (120 V, 1-2 h), and transferred by electroblotting onto nitrocellulose membranes (100 V, 90 min). Proteins were visualised on an Odyssey Fc imager (Licor) using p38 rabbit monoclonal antibody (D13E1, Cell Signaling Technologies #8690, 1:1000), mouse α-tubulin expression (DM1A, Cell Signaling Technologies #3873, 1:1000), and secondary antibodies (Licor anti-rabbit IRDye 680RD #925-6807 or anti-mouse IRDye 800CW #925-32210, 1:10000). Band intensity was quantified using Image Studio Lite (Licor).

### Statistical analysis

All experiments were undertaken in at least three biological replicates, including TCID_50_ infectivity assays with technical quadruplicates and qPCR assays with technical duplicates.

Statistical analysis was performed using GraphPad Prism. Distribution of data was determined by a Shapiro-Wilk test. Significance of data was determined by a two-tailed T test (2 groups) or one-way ANOVA with Tukey’s post-hoc test (>2 groups) for normally distributed data. Where assumptions of normality were violated, significance was determined by a Mann Whitney U (2 groups) or Kruskall-Wallis test with multiple comparisons (>2 groups). Significance levels were defined as *, p≤0.05; **, p≤0.01; ***, p≤0.001.

## Acknowledgements

This work was funded by a Springboard Award to M.S. supported by the Academy of Medical Sciences (AMS), Wellcome Trust, the Government Department for Science, Innovation and Technology (DSIT), the British Heart Foundation and Diabetes UK [SBF009\1124].

---

**Figure S1:**
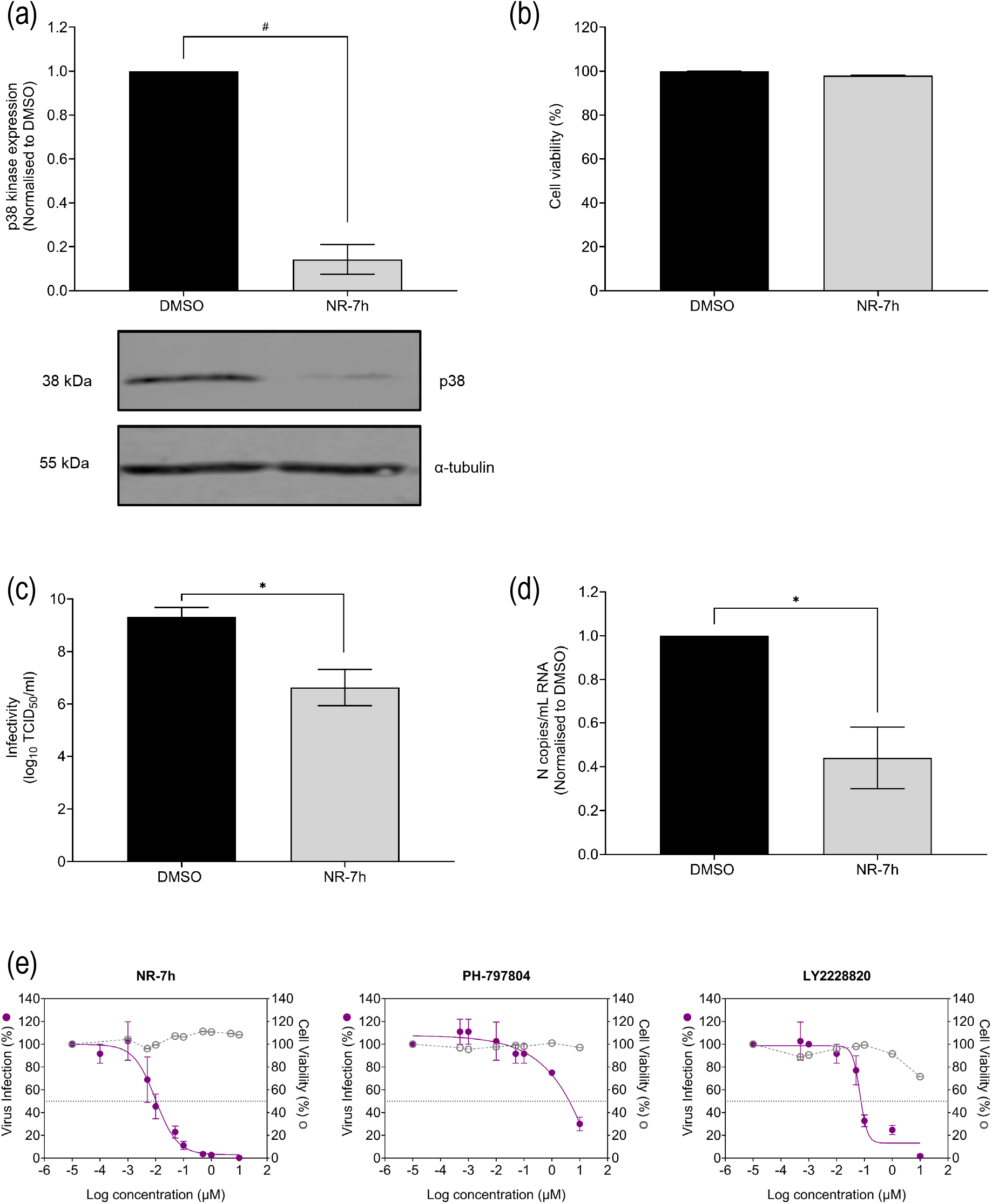
p38α/β-degrading PROTAC inhibits OC43 in BHK-21 cells. BHK-21 cells treated with 10 μM NR-7h for 24 h show significant degradation of p38 kinase **(a)** but no effects on cell viability, as measured by MTS assay **(b)**. BHK-21 cells infected with OC43 (MOI=0.1) and treated with 10 μM NR-7h showed significant reduction of viral infectivity **(c)** and viral RNA **(d)** after 72 h. **(e)** NR-7h showed greater inhibition in BHK-21 cells compared to two p38 kinase small molecule inhibitors (NR-7h, IC_50_ 9.9 nM; PH-797804, IC_50_ 3.5 µM; LY2228820, IC_50_ 69.3 nM). Graphs show mean ± SEM (n=3); *, p≤0.05; **, p≤0.01; ***, p≤0.001. Graphs show mean ± SEM (n=3); *, p≤0.05.

**Figure S2:**
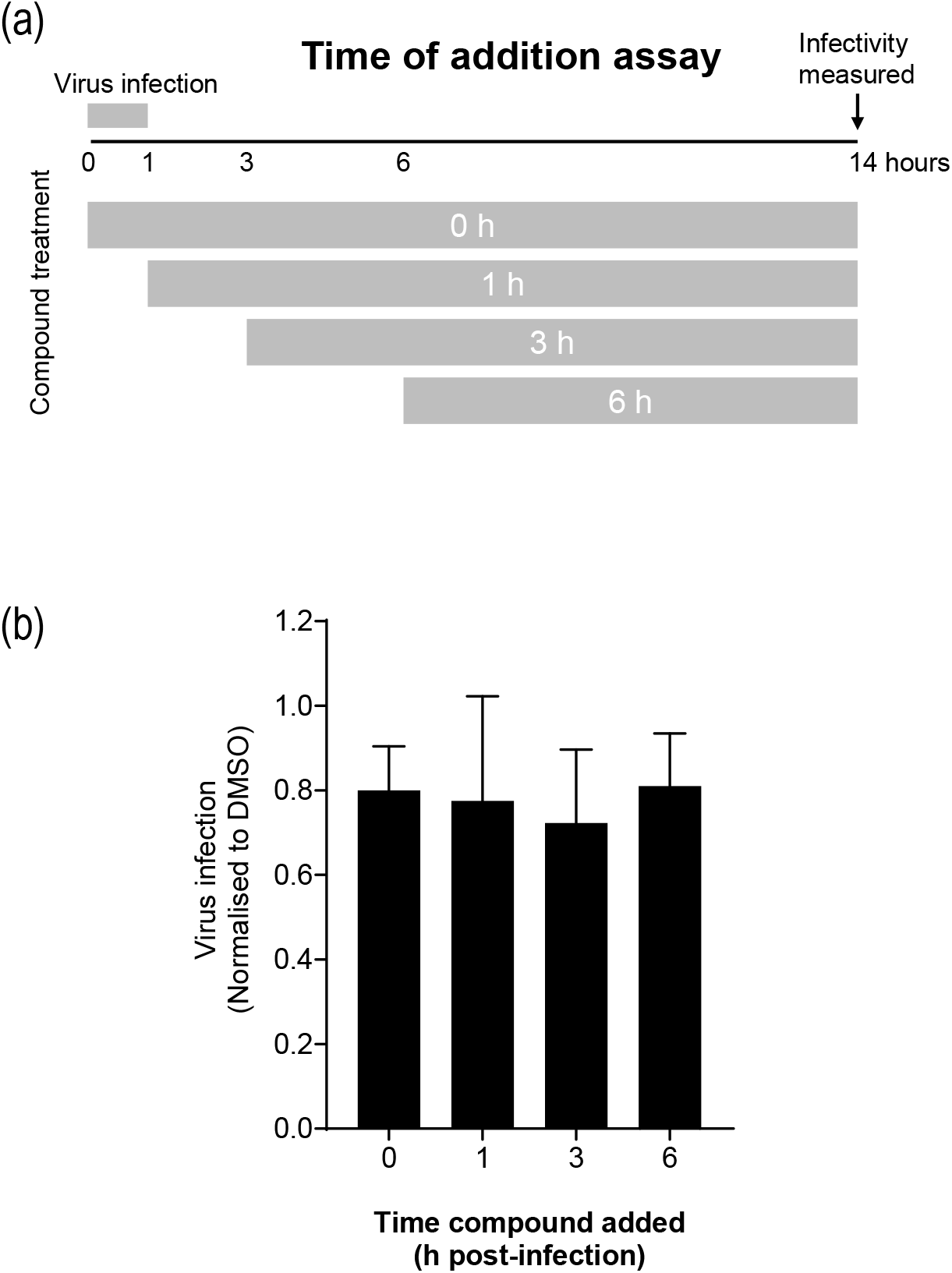
p38 degradation by NR-7h is insufficient within a single infection cycle. **(a)** BHK-21 cells were infected with OC43, and treated with 10 µM NR-7h at 0, 1, 3, or 6 h post-infection, and quantified 14 h post-infection. (b) Quantification of infection was normalised to DMSO-treated cells. Inhibition of infection was not observed for any of these conditions. Graphs show mean ± SEM (n=3).

**Figure S3:**
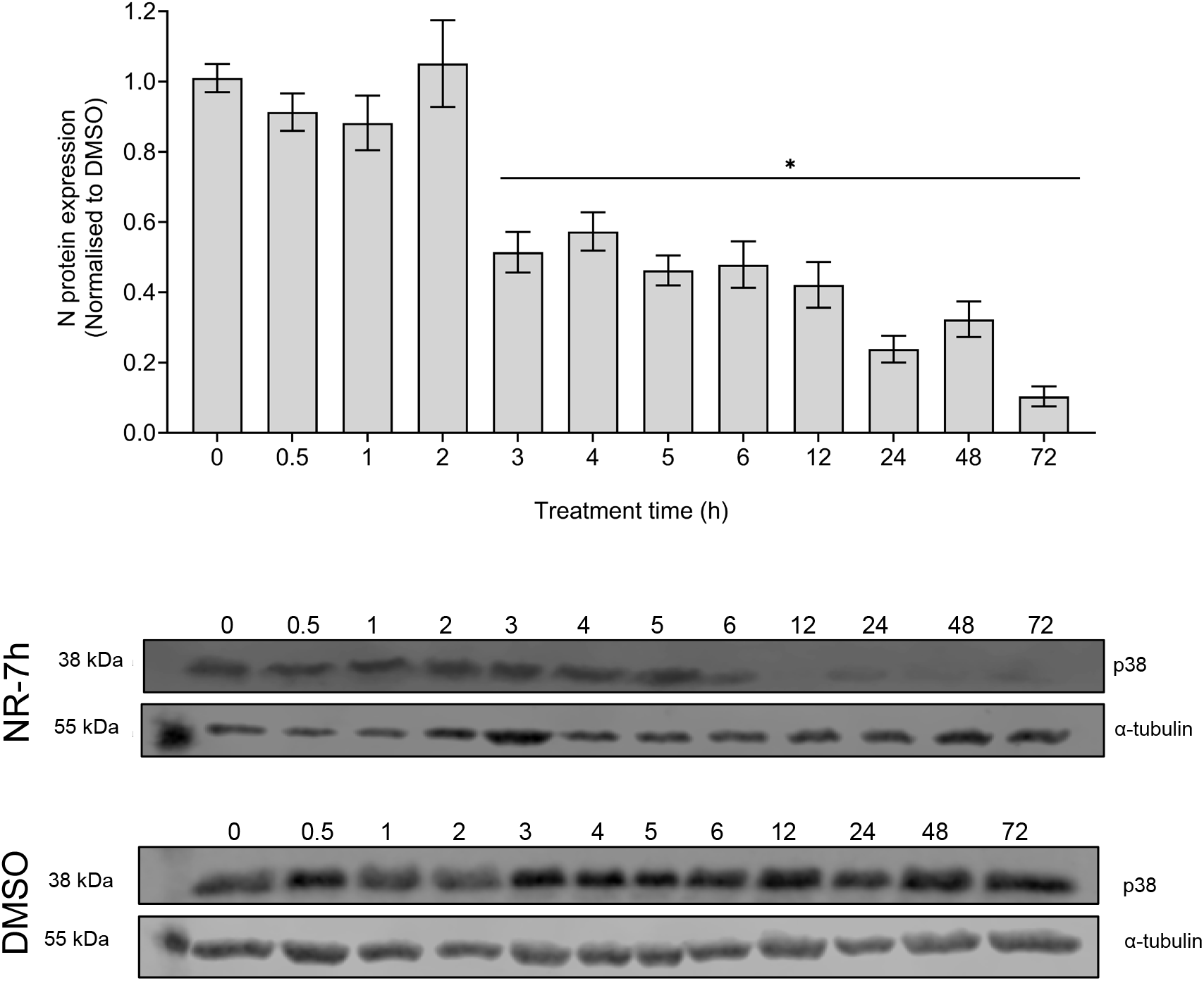
Degradation of p38 kinetics in BHK-21 cells. BHK-21 cells were treated with 10 µM NR-7h or DMSO, and lysates blotted for p38 at various timepoints up to 72 h. p38 was normalised to α-tubulin, then to the DMSO-treated control. DMSO-treated cells showed no degradation of p38, while in NR-7h-treated cells, p38 degradation was observed from 3 h after treatment. Graphs show mean ± SEM (n=3); *, p≤0.05

