## Supplementary Figures S1-3 for "Proof-of-concept of targeted degradation of p38α/β MAPK host-kinase as a potent inhibitor of coronaviruses"

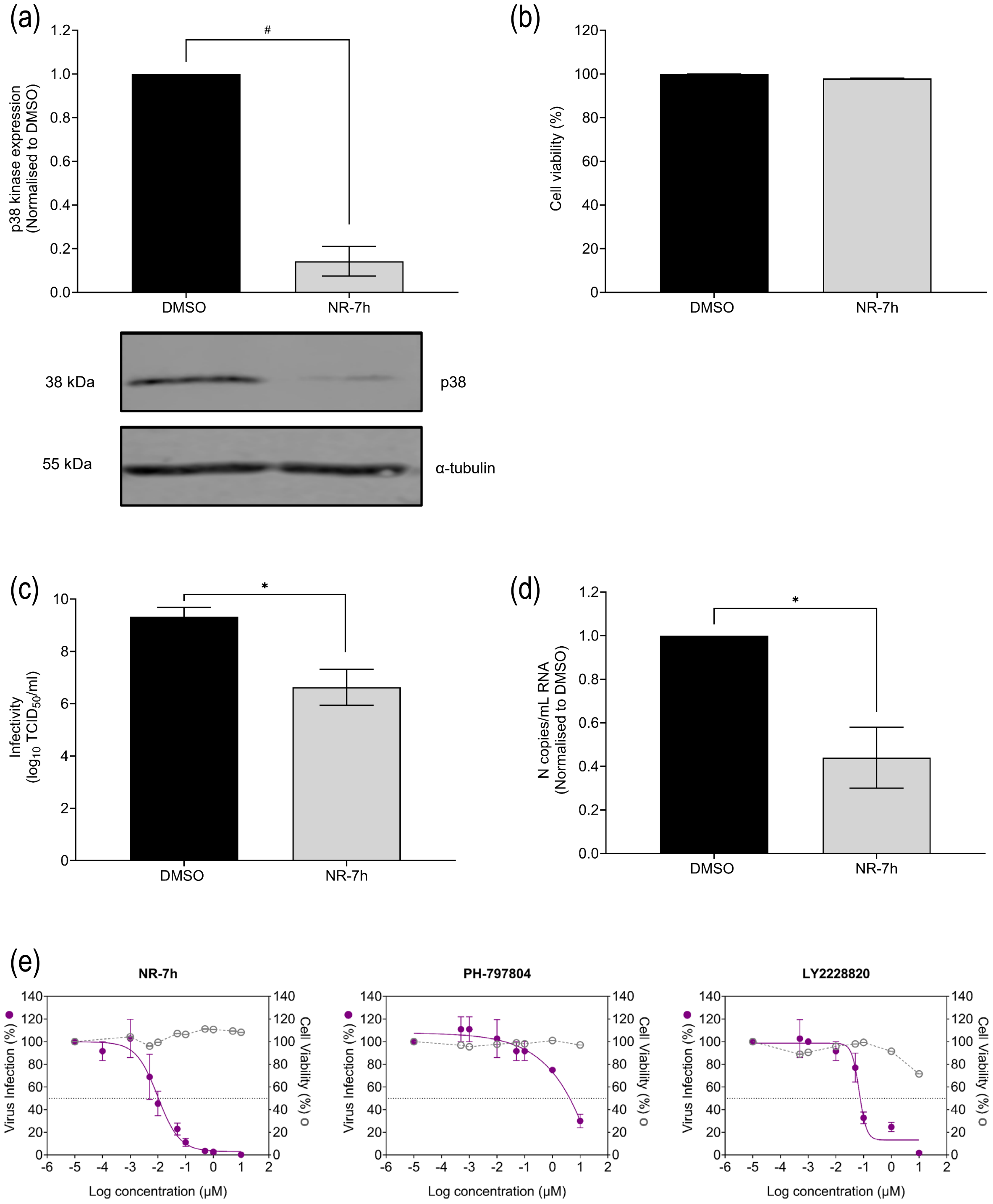


**Figure S1: p38α/β-degrading PROTAC inhibits OC43 in BHK-21 cells.** BHK-21 cells treated with 10 μM NR-7h for 24 h show significant degradation of p38 kinase **(a)** but no effects on cell viability, as measured by MTS assay **(b)**. BHK-21 cells infected with OC43 (MOI=0.1) and treated with 10 μM NR-7h showed significant reduction of viral infectivity **(c)** and viral RNA **(d)** after 72 h. **(e)** NR-7h showed greater inhibition in BHK-21 cells compared to two p38 kinase small molecule inhibitors (NR-7h, $IC_{50}$ 9.9 nM; PH-797804, $IC_{50}$ 3.5 μM; LY2228820, $IC_{50}$ 69.3 nM). Graphs show mean ± SEM (n=3); *, p≤0.05; **, p≤0.01; ***, p≤0.001. Graphs show mean ± SEM (n=3); *, p≤0.05.

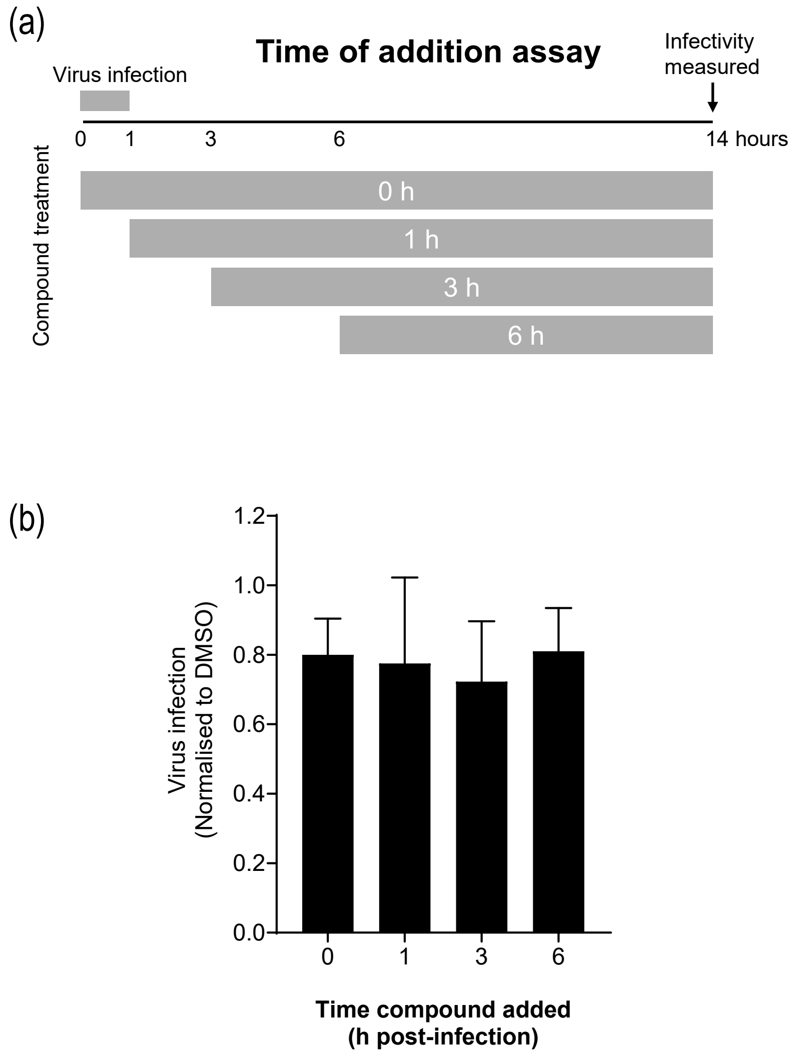


**Figure S2: p38 degradation by NR-7h is insufficient within a single infection cycle.** **(a)** BHK-21 cells were infected with OC43, and treated with 10 µM NR-7h at 0, 1, 3, or 6 h post-infection, and quantified 14 h post-infection. (b) Quantification of infection was normalised to DMSO-treated cells. Inhibition of infection was not observed for any of these conditions. Graphs show mean ± SEM (n=3).

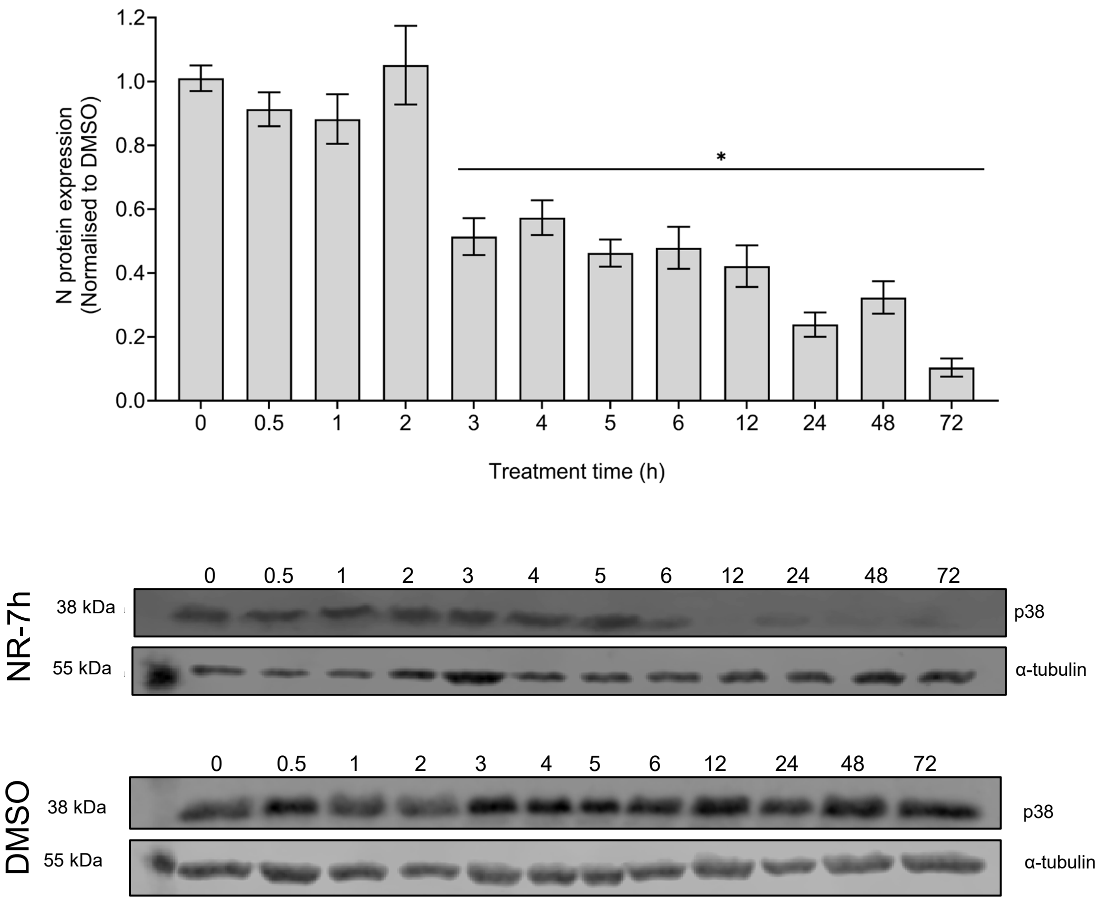


**Figure S3: Degradation of p38 kinetics in BHK-21 cells.** BHK-21 cells were treated with 10 μM NR-7h or DMSO, and lysates blotted for p38 at various timepoints up to 72 h. p38 was normalised to α-tubulin, then to the DMSO-treated control. DMSO-treated cells showed no degradation of p38, while in NR-7h-treated cells, p38 degradation was observed from 3 h after treatment. Graphs show mean ± SEM (n=3); *, $p \leq 0.05$
